# Population-level, state-dependent response as a trait predicting species redistribution under climate change

**DOI:** 10.64898/2026.02.16.706069

**Authors:** Takamitsu Ohigashi, Reiji Masuda, Masayuki Ushio

## Abstract

Species change their population sizes and distributions in response to fluctuating environments. Predicting such changes across space, particularly distributional shifts under climate change, is a central challenge in ecology and conservation^1^. Traditionally, static traits such as habitat preferences, or performance traits such as abundance–environment relationships, have been used to characterize species’ responses to the environment^2,3^. However, these approaches assume fixed relationships between species and their environments, overlooking the inherently state-dependent nature of ecosystems. Here, we show that population-level, state-dependent responses of species to environments can serve as a novel trait to better represent population dynamics in nature, which we term a dynamic response trait. Using nonlinear time-series analysis^4,5^ and long-term marine fish community data collected from Kyoto, Japan^6^, we identify causal influences of water temperature on about one hundred fish species. Species with higher latitudinal centers generally show negative dynamic responses to warming, whereas those with lower latitudinal centers show positive ones. Intriguingly, these dynamic response traits quantified at a single location explain the fish species’ poleward range shift velocities estimated from public biodiversity records; species with negative dynamic responses to warming shift poleward more rapidly, whereas those with positive ones tend to remain. Our findings establish dynamic response traits as a new dimension of trait-based ecology, capturing state-dependent species’ responses. By linking local population dynamics to broad-scale distributional shifts, this approach provides a powerful basis for guiding ecology and conservation under climate change.

## Main

In natural ecosystems, organisms occupy their ecological niches defined by abiotic and biotic conditions; species change their population sizes and distributions in response to environments^7^. Complex relationships between environments and species abundances or community structures have been revealed^8^, yet we still lack a general framework capable of accurately predicting how species and communities will respond to fluctuating environments^9^. In the oceans, many species, including fish, are shifting their distributions poleward under ongoing global warming, often tracking isotherm shifts more clearly than terrestrial species^10,11^. However, the observed shifts are not always consistent with those expected from temperature changes (i.e., not all species move poleward), underscoring the difficulty of predicting species’ range shifts^12–14^. Improving our ability to forecast such ecosystem dynamics remains a critical challenge for biodiversity conservation and resource management under accelerating climate change.

One promising direction is utilizing species traits to predict ecological dynamics in response to fluctuating environments because traits mediate how species interact with the biotic and abiotic environments^2,3,15^. Traditional approaches have relied on static traits—such as morphology or habitat type—assuming that their associated environmental responses remain constant, over a certain time period^2^ (Fig. 1a). For example, fish habitat type and dietary generalism are considered to influence the ability of species to establish in new regions and thus the extent of their range shifts^14^. While these traits are useful for comparisons among categories (e.g., among habitat types), they often ignore the dynamic nature of species’ responses to fluctuating environments, limiting their utility in nature^3,16,17^. As alternatives that capture how species’ performances (e.g., abundance, growth, or survival) change with environmental gradients, performance traits have recently been introduced^3^ (Fig. 1a). These traits are typically quantified as slopes or curves derived from regression-based models, such as linear or generalized additive models (GAM), often measured in laboratory experiments^18^. Compared with static traits, this approach has made progress in trait-based ecological inference, particularly in linking species’ response diversity to community stability to some extent^3^.

**Figure 1.**
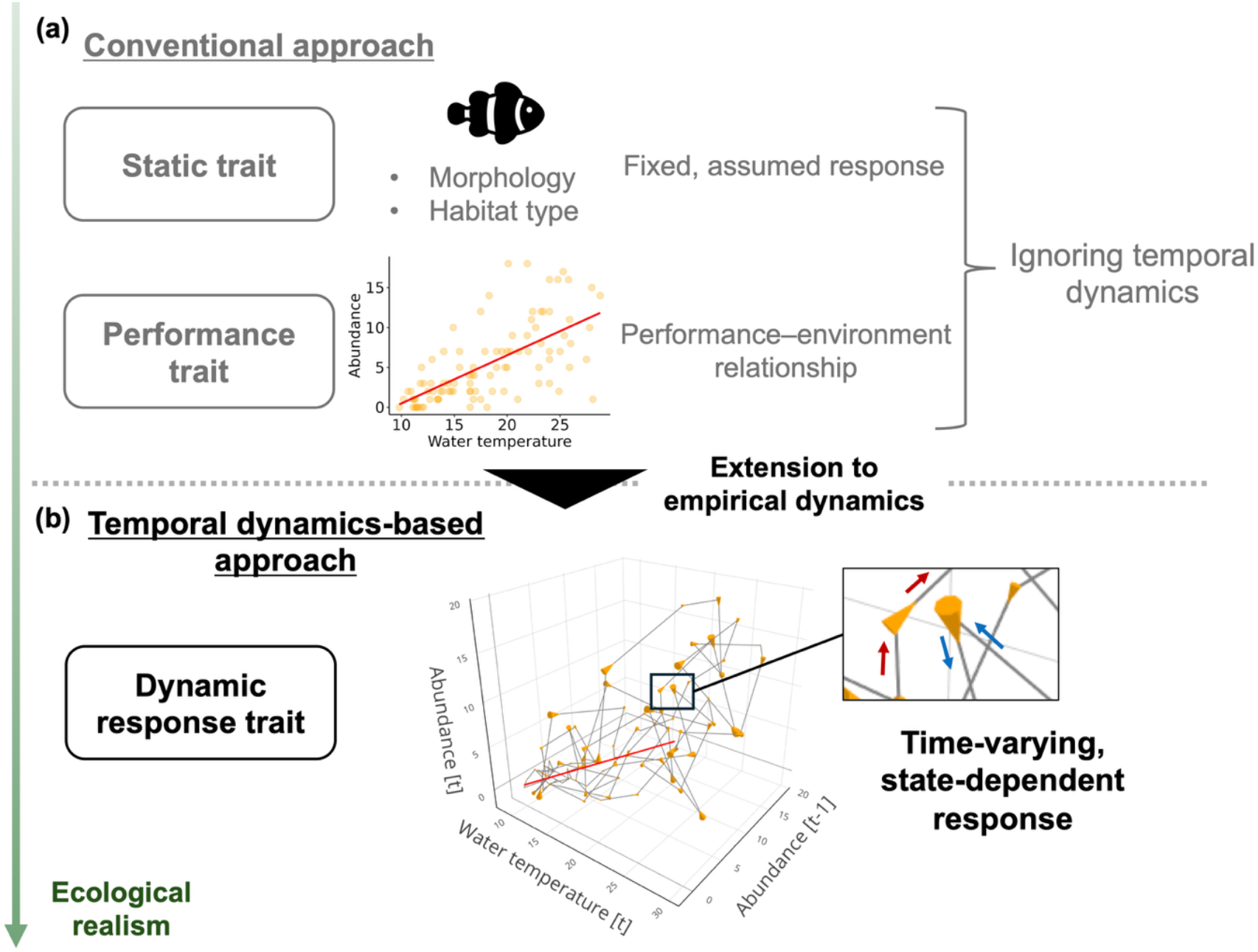
Conceptual diagram illustrating the difference between conventional and dynamic response traits. **a**, Conventional response traits, including both static characteristics (e.g., pelagic lifestyle) and measured performances (e.g., abundance changes along a temperature gradient). **b**, Dynamic response traits, which allow species’ responses to vary depending on the ecological state (e.g., recent population history). Notably, even under the similar environmental conditions, responses can differ. Red and blue arrows indicate such state-dependent dynamics.

In nature, however, even species’ performance–environment relationships depend on complex, multivariate ecological states; responses are inherently nonlinear, shaped by factors such as population density, interspecific interactions, and the recent history of environmental conditions^19,20^. Moreover, species may exhibit different time lags in responding to environmental changes at the population level^19,21^. As a result, correlations between variables evaluated at the same time point may lead to spurious performance–environment relationships (mirage correlations), when the true effects are temporally lagged or even nonexistent^21^. These facts highlight two key limitations of static and performance traits: (1) by abstracting responses— assuming fixed responses or using regression-based slopes or curves—they cannot utilize information about how species act in multivariate states (responses may depend on the present state and/or the past history of biotic/abiotic conditions); and (2) they do not identify which time lags of environmental drivers exert the strongest influence on population responses, which may vary among species (there can be a time lag between population responses and the “causal” multivariate state). Consequently, to better forecast species’ range shifts in a dynamic ecosystem, there is a need to quantify species’ responses in a framework that incorporates present and recent ecological states that reflect the most influential time lags of environmental drivers, thereby capturing “state-dependent” dynamic responses (Fig. 1b).

Such state-dependent dynamic responses of species can be captured by state-space reconstruction (SSR)-based time series analyses, such as empirical dynamic modeling (EDM)^22–24^. By reconstructing population dynamics in a multivariate state space, EDM enables the quantification of species’ responses that depend on ecological states, overcoming the limitations of static or performance traits that assume fixed responses. Moreover, by evaluating responses across multiple time lags, EDM elucidates the most influential species-specific lagged effects of environmental drivers, thereby avoiding spurious correlations^23^. Since such state-dependent responses can reflect intrinsic characteristics in interactions with environmental variability, we argue that they can be interpreted as population-level traits. We therefore propose a new trait index, the “dynamic response trait,” defined as the state-dependent effect of environmental variables on species dynamics that incorporate recent states, at the population level (Fig. 1b).

To demonstrate that the dynamic response trait represents a key ecological property linking species’ environmental sensitivity to climate-driven redistribution, we focus on the temperature responses of fish species. We begin by analyzing long-term (>22 years) underwater visual census data collected twice a month in Maizuru Bay, Kyoto, Japan^6,24^, which include 540 time points of 113 fish species’ abundances and water temperature data (Fig. 2a). In brief, the fish communities exhibited clear seasonality and gradual compositional changes across years (Fig. 2b). Over the 22 years, species with low latitudinal distribution centers increased in their abundance and richness, while species with high latitudinal distribution centers decreased, highlighting a shift in latitudinal signatures of the fish communities (Fig. 2c, d).

**Figure 2.**
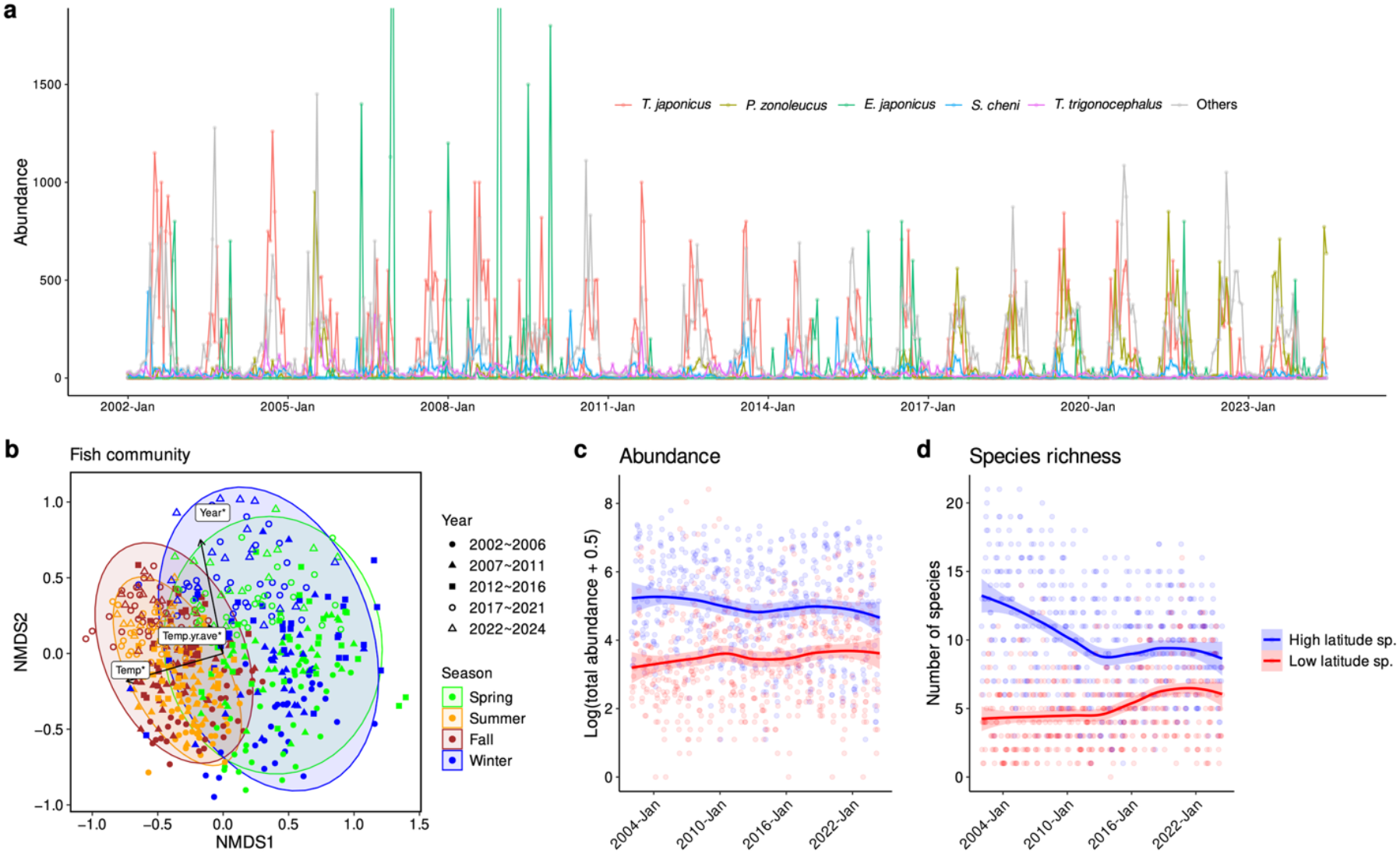
Characteristics of the long-term fish observational data in Maizuru Bay. **a**, Time series of population abundances for the top five most dominant species, along with the summed abundance of the remaining 108 species. For visualization purposes, two extreme values (>1800 for *E. japonicus*) are omitted from the plot. **b**, Seasonal and interannual variation in fish community structure. Non-metric multidimensional scaling (NMDS) is used to visualize community dissimilarities. Environmental variables clearly associated with the community variation are represented as arrows (*p* < 0.05, Temp: water temperature; Temp.yr.ave: annual mean temperature; Year: sampling year). **c–d**, Fish community dynamics grouped by latitudinal distribution category. Species are categorized as “High latitude” or “Low latitude” based on whether their latitudinal distribution center is above or below the median of the 113 observed species. Total abundance (**c**) and species richness (**d**) are shown for each latitudinal category.

Using this dataset, we detect causal relationships between water temperature and fish species using the Unified Information-theoretic Causality (UIC), an EDM method integrating information theory^4^, which quantifies transfer entropy (TE) between temperature and fish species’ time series. By quantifying TE across time lags using the time series, UIC allows us to identify the most influential temperature time lag for each species (see Methods). In our analysis, when assessing the presence of causality, some species were present only during specific intervals within the 22-year monitoring timeframe in Maizuru Bay (e.g., due to range shifts), such that applying UIC to the full series may falsely suggest no causal link with temperature, even when one exists. We therefore apply UIC in 5-year moving windows (120 time points) and test whether temperature has causal effects on fish species in at least one 5-year window. Overall, temperature exhibits statistically clear causal effects on 102 fish species in at least one window (Methods, Extended Data Fig. 1).

For the 102 fish species, we quantify time-varying interaction strengths of water temperature (i.e., the dynamic response of fish species) using the multiview-distance regularized S-map^5^ (MDR S-map, see Methods). The MDR S-map is a weighted local linear regression using low-dimensional SSRs called “multiviews,” which escapes the curse of dimensionality^25^. We perform the MDR S-map using the optimal time lag identified by the UIC analysis for each fish species. This resulted in 56 species that possess ≥1 optimal embedding dimension and meaningful prediction skills (ρ > 0), and we utilize the S-map coefficients as quantitative estimates of their dynamic responses. A positive response indicates that a species’ population size increases with rising temperature, whereas a negative response indicates that it decreases with warming. The fish species’ mean dynamic responses to temperature are clearly negatively correlated with their latitudinal distribution centers, indicating that northern species respond negatively to temperature, whereas subtropical species respond positively (Fig. 3). This relationship persists even after accounting for fish phylogeny using phylogenetic independent contrasts (Extended Data Fig. 2). In contrast, performance traits quantified with simple linear regression or GAM do not show such clear relationships (Extended Data Fig. 3). These conventional performance traits reflect species abundance at a given temperature or a given time point, but disregard the multivariate nature of ecological states, such as recent population trends (e.g., whether they are increasing or decreasing), and time-lagged responses to temperature. This oversight, together with the general tendency for species abundance to increase during warm periods (Extended Data Fig. 6)^19^, can result in unreliable, ecologically less meaningful “trait” estimation. In contrast, dynamic response traits can overcome this limitation by capturing the state-dependent component of temperature effects, reasonably corresponding to their distribution centers.

**Figure 3.**
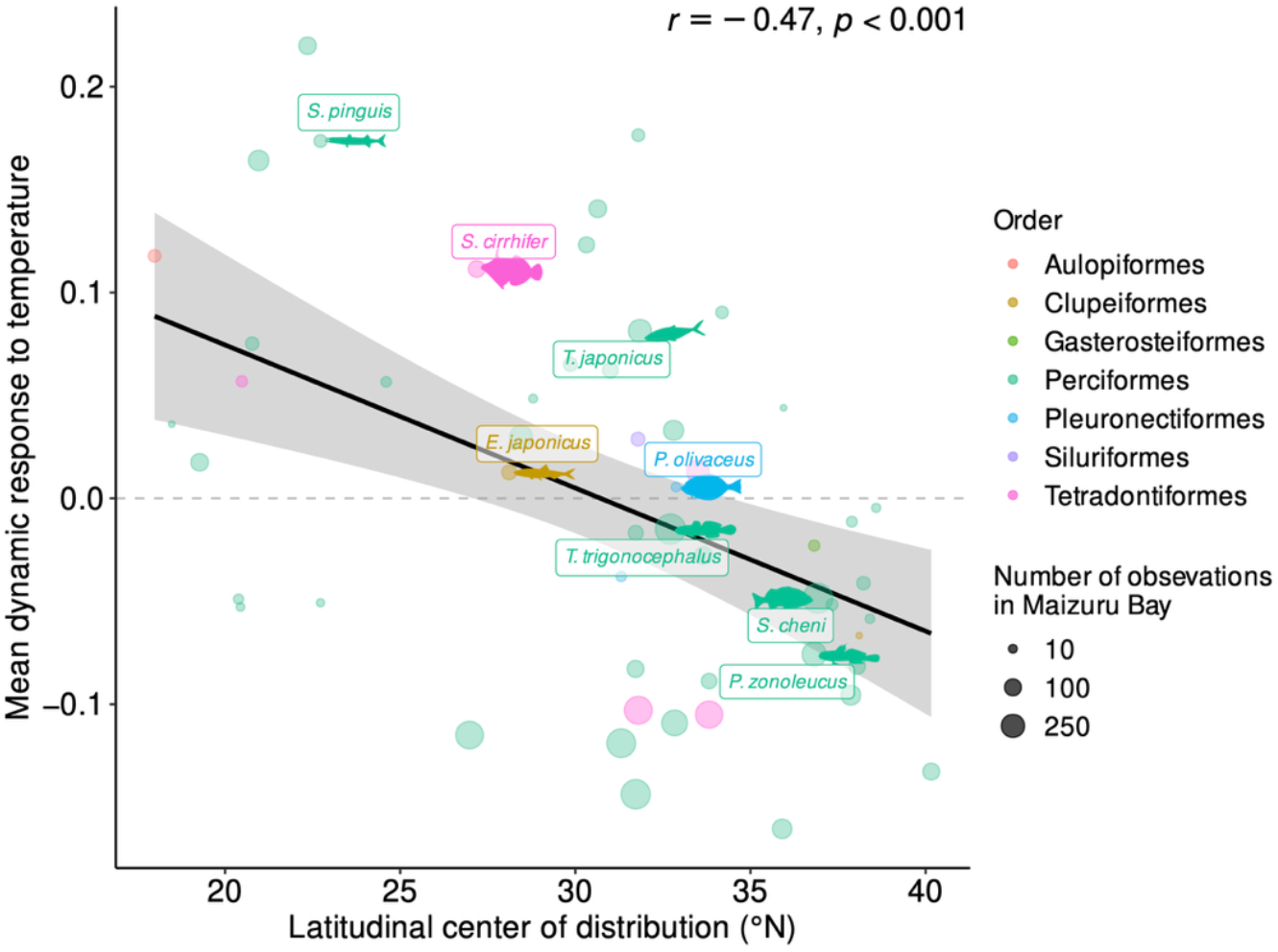
Latitudinal distribution and dynamic responses to temperature of species. The relationship between latitudinal distribution centers and mean dynamic responses to temperature is shown (based on 56 species for which dynamic response to temperature is quantified; two outliers are excluded). Point size represents the number of observations (abundance > 0) in Maizuru Bay during this study. Fish silhouettes and names are shown next to points for the five most abundant species and those classified as “highly commercial” in the FishBase database. These silhouettes are obtained from PhyloPic; detailed credits are provided in the Supplementary Information Section 5. Pearson’s correlation coefficient (*r*) and its *p* value are indicated in the figure.

The negative relationship between species’ dynamic responses to temperature and latitudinal distribution centers suggests that, under continued warming, high-latitude species are likely to decrease in Maizuru Bay, whereas low-latitude species will increase, consistent with long-term changes already observed in the bay (Fig. 2c, d). This may indicate that dynamic response traits capture one of the essential, yet previously overlooked traits of fish species, suggesting these traits have strong potential for predicting species’ range shifts even in other regions.

To independently and further evaluate the performance of the new trait index, we then estimated the poleward range shift velocities of the fish species observed in Maizuru Bay, using public biodiversity data from iNaturalist and GBIF (Methods). Intriguingly, the poleward shift velocities estimated from the public datasets are explained by species’ mean dynamic responses to temperature quantified using the time series in Maizuru Bay (Fig. 4a). This indicates that species showing negative responses to temperature shift poleward more rapidly, whereas those with positive responses tend to persist within their ranges.

**Figure 4.**
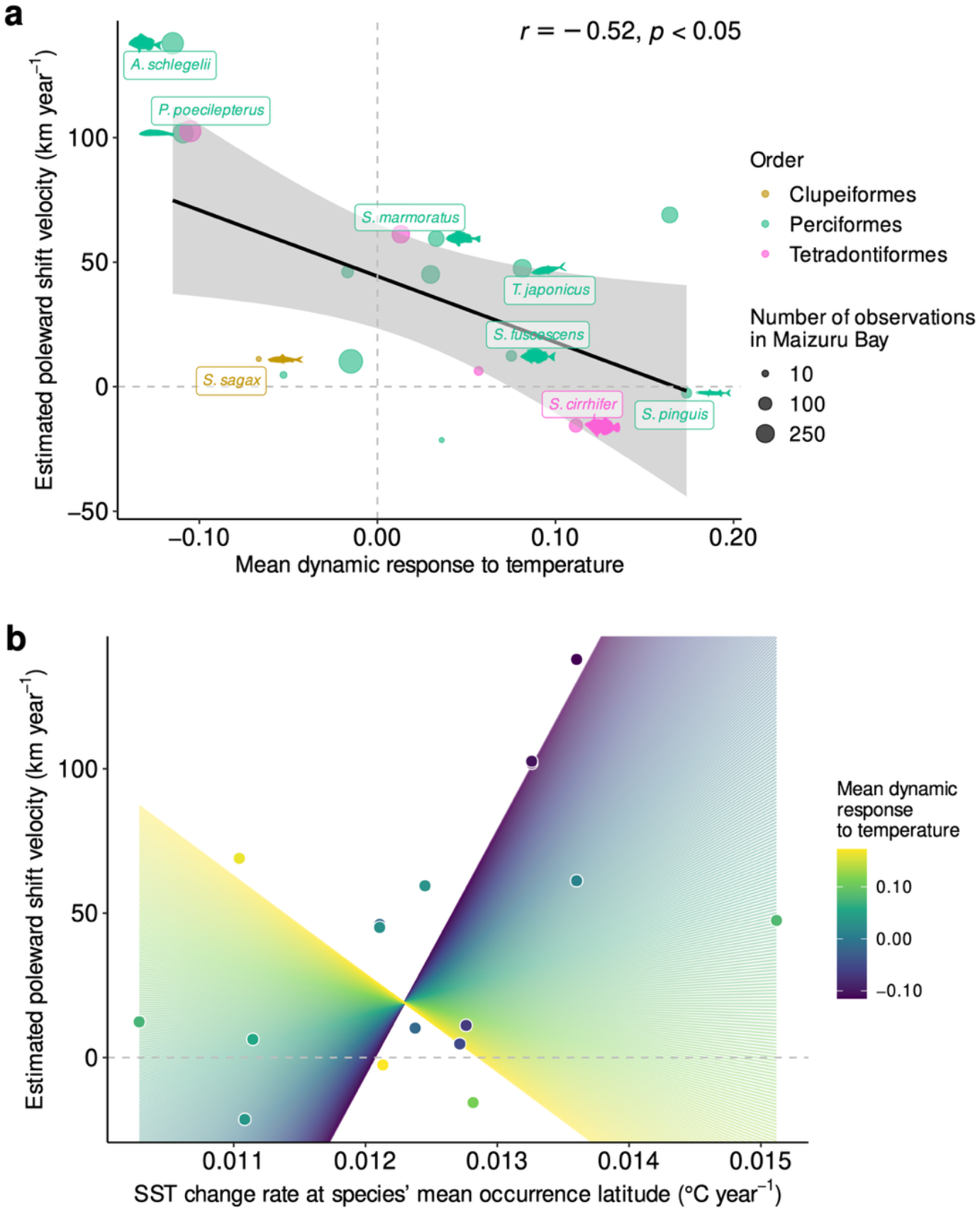
Fish range shift velocity and dynamic response to temperature. **a**, The relationship between each species’ mean dynamic response to temperature and its estimated poleward shift velocity, based on iNaturalist and GBIF occurrence records collected in East Asia and Oceania. Point size represents the number of observations (abundance > 0) in Maizuru Bay during this study. Fish silhouettes are shown for the species classified as “highly commercial” or “commercial” in the FishBase database. These silhouettes are obtained from PhyloPic; detailed credits are provided in the Supplementary Information Section 5. Pearson’s correlation coefficient (*r*) and its *p* value are shown. **b**, Interaction effect of the sea surface temperature (SST) change rate at each species’ mean occurrence latitude (based on iNaturalist and GBIF records) and the mean dynamic response to temperature, on the estimated poleward shift velocity. Colored lines indicate model-predicted relationships between the SST change rate and poleward shift velocity, varying with the mean dynamic response to temperature. Only 17 species, for which both dynamic response to temperature and poleward shift velocity could be quantified, are shown in this figure.

To clarify whether these species’ behaviors are driven by their inherent traits responding to rising temperature, we further investigate the relationships among dynamic response traits, changes in sea surface temperature (SST) experienced by the species (i.e., SST change rate at the mean latitude where the species occurrences were recorded in iNaturalist and GBIF), and the estimated range shift velocities (Fig. 4b). The species with negative mean dynamic responses shift poleward rapidly with increasing SST, whereas those with positive responses tend to remain despite warming (Fig. 4b). This result is consistent with the expectation that species with narrower thermal tolerances contract their ranges at the trailing edge, thereby shifting their range centroids poleward^14^. Simultaneously, the negative response to temperature—which, in turn, increases abundance as temperature decreases—suggests an ability to maintain positive population growth in the higher-latitude, cooler environments of the leading edge^14^. It is widely recognized that conventional thermal limits measured under experimental conditions often fail to reflect realized thermal limits in nature^26,27^, and directly linking contemporaneous species distributions to climatic conditions can obscure the ecological processes underlying redistribution^28^. Thus, predicting species’ distributions requires integrating explicit ecological processes rather than relying solely on correlative approaches^28^. From these perspectives, our dynamic response trait provides a distribution-independent, causal (i.e., process-based) measure of temperature response, serving as a predictor of broad-scale redistribution that accounts for the nonlinear dynamics inherent in natural systems.

Our findings establish the dynamic response trait as a novel trait dimension, extending trait-based ecology to capture the state-dependent responses of species to the environment. This framework is not limited to temperature; in principle, any time series (e.g., pH, oxygen, nutrients) could be used to quantify dynamic response traits, widely applicable across ecological contexts.

One may be concerned that interspecific interactions could bias the estimation of dynamic responses to temperature. Indeed, including other fish species’ time series in the embedding— thereby quantifying the temperature response by excluding interspecific interactions—produces slightly stronger range-shift predictions than using temperature alone, yet the overall pattern remains unchanged (Extended Data Fig. 7, Supplementary Information Section 1). Because obtaining complete multispecies time series that capture all interacting taxa is rarely possible in practice, this consistency is particularly important.

Notably, even though these traits were quantified from time series at a single site, they explain broad-scale distributional shifts, demonstrating their potential in understanding and predicting future changes in biodiversity—a fundamental element of sustainable human societies. While our census data rely on labor-intensive underwater observations, emerging approaches such as environmental DNA and expanding citizen-science surveys can scale up the estimation of dynamic response traits across space, taxa, and crucially, provide the time-series data essential for EDM^22,29,30^. Our study offers a proof of concept, showing that the dynamic response trait captures an important ecological aspect and can also serve as a basis for predicting changes in biodiversity patterns under climate change.

## Methods

### Fish community and temperature time-series data

Long-term time-series data of the fish community and bottom water temperature were collected by underwater observation twice a month along the coast of the Maizuru Fishery Research Station of Kyoto University (Nagahama, Kyoto, Japan: 35° 29′ 24.66″ N, 135° 22′ 5.76″ E) from 1 January 2002 to 20 June 2024, yielding 540 time points over more than 22 years^6,24^. The study area was within 50 m of the shore at depths of 1–10 m. A total 600-m visual transect line was set in three areas: each segment was 200 m long and 2 m wide. Additional details such as vegetation and salinity trends were documented in a previous study^24^.

The census was carried out by scuba diving. Fish species and their abundances observed within 1 m of the transect line were recorded. Observations were conducted on sunny days and commenced around noon during high tide for 2–3 hours. Bottom water temperature was measured at the deepest point of the transect line (10-m depth) during the dive. Visibility ranged from 1 to 15 m but was typically 3–5 m. Importantly, the same scientist consistently led the field surveys throughout the 22-year research period. Thus, inconsistency in fish identification and counting was minimal in the time series. In total, 113 fish species were observed during the period. The latitudinal distribution centers of these fish species were identified based on published information for Japanese fishes^31,32^.

### The Unified information-theoretic causality method

We detected causal interactions between water temperature and fish species using the unified information-theoretic causality (UIC)^4^, a time-series analysis-based causality detection method. UIC integrates two existing approaches for detecting causality in time-series data—transfer entropy^33^ (TE) and convergent cross mapping^22^ (CCM). Consider two variables, an effect variable ***x*** = {*x*_*t*_} and a potential causal variable ***y*** = {*y*_*t*_}, where each represents an observed sequence over time (e.g., ***x*** for the population of a fish species and ***y*** for water temperature). In conventional TE, the information flow from ***y*** to ***x*** is quantified by evaluating whether the entropy of *x*_*t*_ conditioned on *y*_*t*+*tp*_ and ***x*** is greater than that conditioned on ***x*** alone, where *t* is the target time index and *tp* is time step (if *tp* = −1, TE evaluates the causal effect of *y*_*t*−1_ on *x*_*t*_). In CCM, the state space of ***x*** is first reconstructed using time-delay embedding, and *y*_*t*+*tp*_ is predicted (i.e., cross mapping). Then, causality was inferred when the prediction skill improves as the amount of training data increases (i.e., “convergence”). UIC combines the two approaches: it quantifies information flow between variables as TE through cross mapping^4^.

In UIC, similarly to other EDM approaches, the optimal number of dimensions for the time-delay embedding (*E*) is selected based on the prediction skill of the local regression (simplex projection) using a state space including the potential causal variable, ***y***, and the effect variable, ***x***. In UIC, this is implemented using the framework of TE as follows:

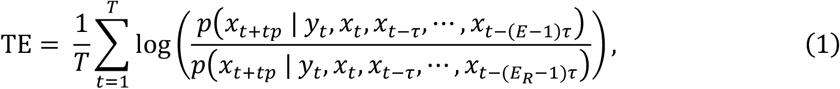

where *T* is the total number of points in the reconstructed state space (the total number of time points − *E* + 1); *E*_*R*_(< *E*) is the optimal embedding dimension for lower-dimensional models; *t* is the time index; *tp* is the time step; and τ is the time lag unit. *p*(*A*|*B*) denotes the conditional probability of *A* given *B*. Equation (1) is a TE-based version of the simplex projection method, and when *tp* = 1, *E* is determined by the value that maximizes TE on one-step-ahead prediction. Then, with the optimal embedding dimension *E*, UIC quantifies information flow from ***y*** to ***x*** in the following way:

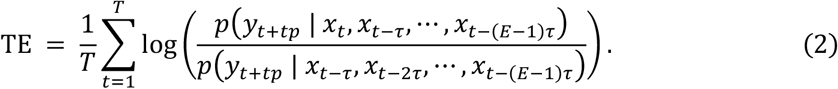

For example, if *tp* = −1 in Equation (2), UIC evaluates the causal effect of *y*_*t*−1_on *x*_*t*_.

In the present study, we tested causality with time lags (i.e., *tp* in Equation (2)) up to −12, corresponding to a 6-month time lag, using the “rUIC” package^34^ (v0.9.12). To evaluate the statistical clarity of TE, we generated 1,000 surrogate water temperature time series using the “seasonal” method implemented in the make_surrogate_data function in the “rEDM” package^35^ (v0.7.5), which preserves seasonal structure while randomizing anomalies. After standardizing temperature and species abundance (mean = 0, SD = 1), TE from each surrogate temperature to the target species was calculated, and the *p*-value was estimated as the proportion of surrogate-based TE values that exceeded the observed TE at each time lag. To account for multiple comparisons across species, the resulting *p*-values were adjusted at each time lag using the Benjamini–Hochberg false discovery rate correction across all 113 species.

While the initial UIC analysis was performed using the full 540-timepoint dataset, this approach may miss causal relationships for species whose occurrence in Maizuru Bay is temporally restricted—e.g., species that arrived later or disappeared over time due to range shifts. To address this, we conducted a moving-window UIC approach using a window size of 120 timepoints (5 years), sliding across the entire time series (i.e., 421 windows in total). For each window, we applied the same TE calculation and surrogate testing procedure as described above. A species was considered to exhibit causal influence from temperature if at least one window yielded a statistically clear TE (BH-adjusted *p* < 0.05, adjusted across all windows and species). This moving-window UIC analysis identified 102 species with causality at least one window, and these species were included in the subsequent analysis, the MDR S-map.

### The MDR S-map method

We quantified the species’ dynamic responses to temperature (that is, also called S-map coefficients, or interaction strengths particularly in studies on interspecific interactions^19^). To estimate the effects of interacting variables on the future dynamics of a target variable, we used an improved version of the sequential locally weighted global linear map (S-map)^36^, known as the multiview distance regularized S-map (MDR S-map)^5^. Consider a system consisting of *E* interacting variables (including possible time-delay coordinates), where the state of the system at time *t* is represented as ***x***_***t***_ = {*x*_1,*t*_,*x*_2,*t*,_ ⋯, *x*_*E,t*_}F. For each target time point *t*, the S-map constructs a local linear model to predict the future value *x*_1,*t*+*tp*_ using the reconstructed state space vector ***x***_***t***_. That is,

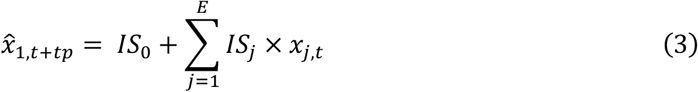

where 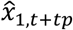 is the predicted value of *x*_1_ at time *t* + *tp*; *IS*_*j*_ is the regression coefficient that can be interpreted as the interaction strength; and *IS*_0_ is the intercept of the linear model. The linear model is fitted using the remaining vectors in the state space, with greater weights assigned to points closer to the target vector ***x***_***t***_ (i.e., locally weighted linear regression). Equation (3) is based on Takens’ embedding theorem for univariate systems^37^, which can be generalized to multivariate systems^38^. In our study, we extended the S-map to a multivariate form by including the water temperature at the time lag that showed the strongest causal effect (i.e., the largest TE) on the fish species according to the UIC analysis as:

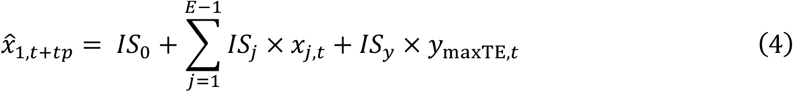

where *y*_*maxTE,t*_ is the time-delayed y (e.g., water temperature) with the largest TE on 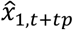 (e.g., future value of a fish population) at *t*; and *IS*_*y*_ is the time-varying interaction strength between *y*_*maxTE,t*_ and 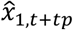. We consider *IS*_*y*_ in Equation (4) as a state-dependent, population-level response to water temperature for a fish species, referred to as a “dynamic response,” which may reflect a trait-like characteristic of the species, although we acknowledge that a trait should be defined at the individual level^2^.

In a typical S-map, the distances between points are typically measured using Euclidean distance. However, in high-dimensional state spaces, Euclidean distance often fails to identify true nearest neighbors. The MDR S-map addresses this limitation by replacing Euclidean distance with the multiview distance^5^, which is computed by ensembling distances across multiple low-dimensional projections of the original state space (i.e., multiview embedding^25^). Additionally, to reduce the possibility of overestimation and improve forecasting performance, the MDR S-map incorporates regularization^39^ (i.e., ridge regression). This method quantifies a species’ dynamic response to temperature at each time point, allowing for time-varying estimates (i.e., the maximum number of dynamic response values is equal to the number of time points − the optimal embedding dimension + 1). All MDR S-map analyses were conducted using our custom R package “macam”^40^, after standardizing the temperature and species abundances as in the UIC. Although interaction strengths are often theoretically defined on a per-capita basis^41^, species’ range shifts represent population-level processes rather than per-capita demographic responses; thus, we focus on population-level dynamic responses as the primary quantity of interest in this study.

### Estimation of species range shift velocity

To test whether species’ dynamic response traits are capable of explaining their relatively long-term, broad-scale range shifts under climate change, we investigated the relationship between dynamic responses and range shift velocities. We utilized public biodiversity data from iNaturalist and the Global Biodiversity Information Facility (GBIF) to estimate the range shift velocities for the fish species observed in our study site (Maizuru Bay). The “rinat” and “rgbif” packages^42,43^ were used to retrieve public occurrence data from 1960 to 2024, with a maximum of 10,000 records per species per year. Data containing terms such as “aquarium,” “zoo,” “museum,” and “market” in Japanese, English, Spanish, French, Chinese, and Korean were excluded, as these observations were from artificial environments. Additionally, we used the “CoordinateCleaner” package^44^ to remove records from capital cities, country centroids, and duplicated data points.

Species with at least 10 records in each year, spanning at least 5 years, from observations covering East Asia and Oceania (i.e., any latitude within 105–180°E) were included in the estimation of shift velocities. To avoid over-reliance on extreme records for the calculation of shift velocity, we excluded the data points within the upper and lower 5% of absolute latitudes for each species in each year^45^. Then, to account for sampling bias due to observation frequency, we calculated observation effort as the total number of records including all fish species per degree of latitude per year. Finally, species-specific poleward shift velocities were estimated as the slope coefficients from linear regressions while accounting for the observation efforts at each latitude band (i.e., *lm*(*abs*(*latitude*)∼*year, weight* = 1/*effort*) in R). Additionally, we compared the estimated shift velocities with long-term abundance trends at Maizuru Bay to assess the consistency between them (Extended Data Fig. 8, Supplementary Information Section 2).

### Sea surface temperature change

To examine whether the estimated range shifts were consistent with the species’ dynamic response traits and the extent of the temperature change they experienced, we quantified the long-term trends in sea surface temperature (SST) at the latitudinal ranges inhabited by each fish species observed in Maizuru Bay. Specifically, we used the HadISST1 dataset^46^ to extract SST data for the region between 105°E and 180°E, covering East Asia and Oceania. For each species, we identified the mean observed latitude based on occurrence records from iNaturalist and GBIF, using the latitude values incorporated in the estimation of the species range shift velocity described in the previous section. If a species occurred in both hemispheres, we separately calculated the SST trend (in °C per year) at its mean latitude in each hemisphere using a linear regression over the period 1960–2024 with the lm function, and then averaged the two values. If a species was found in only one hemisphere, we used the SST trend from that hemisphere only.

### Correlation and regression analyses

We tested the correlations between species’ latitudinal distribution centers, range shift velocities, and their dynamic responses to temperature using Pearson’s correlation tests with the cor.test function in R. To account for phylogenetic influences on these correlations, we additionally performed phylogenetic independent contrasts (PIC) analyses (Extended Data Fig. 2, 3c–d, 4, 5c– d, Supplementary Information Section 2). To perform the PIC analyses, we first collected mitochondrial and protein-coding genomes available in public databases (MitoFish^47^ and NCBI) for the 113 species observed in Maizuru Bay. DNA sequences were aligned for each gene using the ClustalW^48^ via the “msa” package^49^ and concatenated into a supermatrix using the “phytools” package^50^. A phylogenetic tree was then constructed with RAxML^51^ using the raxml function in the “ips” package^52^, while constraining the tree topology to a family-level bony fish backbone tree based on a previous study^53^. Finally, we computed PICs using the pic function in the “ape” package^54^ and assessed correlations among the resulting values.

To evaluate the effects of species’ dynamic responses to temperature and local SST changes on range shift velocity, we fitted a linear model including an interaction term between these two explanatory variables, using the lm function.

## Author contributions

TO and MU conceptualized the study and developed the methodology. TO performed the data analysis and drafted the manuscript. RM conducted the long-term field surveys and contributed to the ecological interpretation of the results. All authors reviewed and approved the final manuscript.

## Acknowledgements

We thank Masaki Miya for providing advice on the phylogenetic analysis, and Chih-hao Hsieh for discussions regarding the ecological applications of time-series analysis. This research was supported by JSPS KAKENHI Grant Number 24KJ0018 to TO, and The Hong Kong University of Science and Technology Startup Fund to MU.

## Code availability

The R scripts used for the main analyses are available at GitHub (https://github.com/taka-ohi/DynaResp_Maizuru_Fish).

## Data availability

Time-series data of the Maizuru fish community are available at GitHub (https://github.com/taka-ohi/DynaResp_Maizuru_Fish). All other data are available from the corresponding author(s) upon reasonable request.

## Conflict of interest statements

The authors declare no conflicts of interest.

## Notes

### Competing Interest Statement

The authors have declared no competing interest.

