## Extended Data for "Population-level, state-dependent response as a trait predicting species redistribution under climate change"

### Contents:

- **Extended Data Figure 1**| Moving-window analysis of causal effects of temperature on fish species dynamics
- **Extended Data Figure 2**| Relationship between latitudinal distribution and dynamic response to temperature accounting for phylogeny
- **Extended Data Figure 3**| Relationships between latitudinal distribution and conventional temperature response traits
- **Extended Data Figure 4**| Relationship between fish range shift velocity and dynamic response to temperature accounting for phylogeny
- **Extended Data Figure 5**| Relationships between conventional temperature response traits and fish range shift velocities
- **Extended Data Figure 6**| Relationships between water temperature and fish communities grouped by latitudinal category
- **Extended Data Figure 7**| Dynamic responses to temperature after excluding interspecific interactions
- **Extended Data Figure 8**| Relationships between species' latitudinal distribution centers and their estimated poleward shift velocities, with temporal changes in local abundance

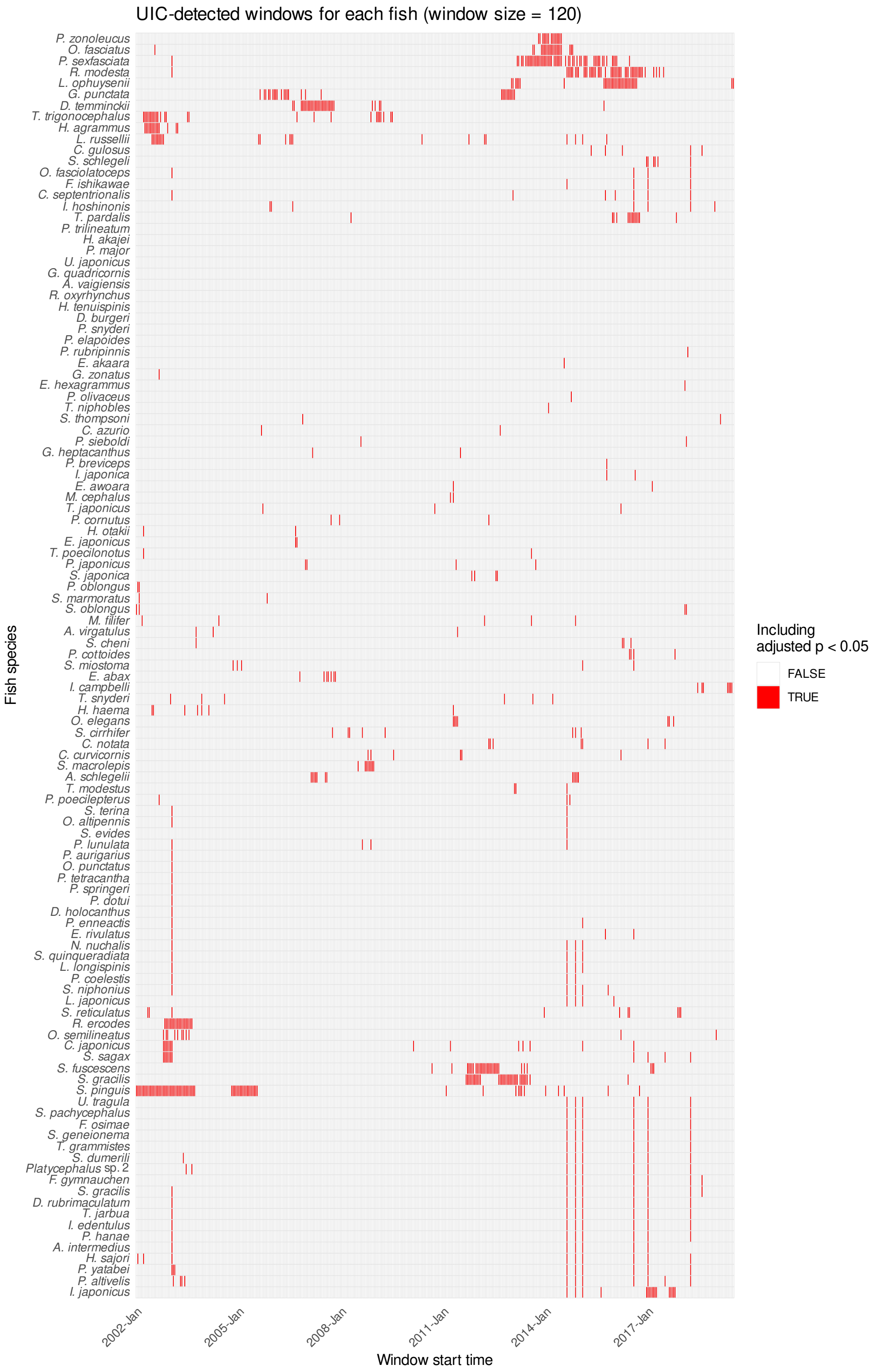


**Extended Data Figure 1| Moving-window analysis of causal effects of temperature on fish species dynamics.** Results of the moving-window Unified Information-theoretic Causality (UIC) analysis conducted with a five-year window (120 time points) to detect causal influences from temperature to each fish species. The heatmap shows window-specific significance, where the horizontal axis indicates the window start time and the vertical axis represents fish species. Windows with BH-adjusted $p < 0.05$ are shown in red. Species are ordered based on hierarchical clustering (Ward.D2 linkage on Euclidean distance) using the binary significance matrix.

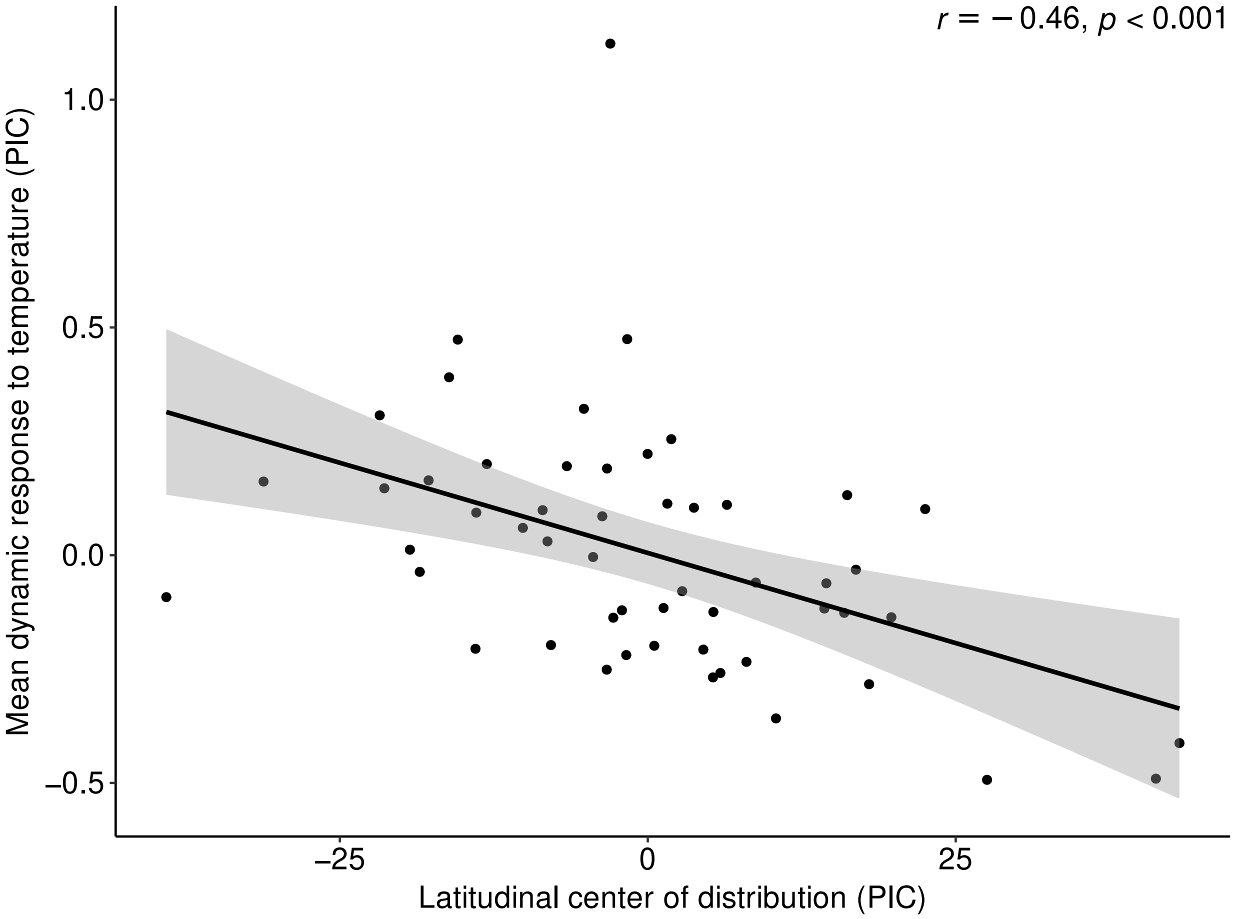


**Extended Data Figure 2| Relationship between latitudinal distribution and dynamic response to temperature accounting for phylogeny.** The relationship between species' latitudinal distribution centers and their mean dynamic responses to temperature is shown, accounting for phylogenetic relatedness using phylogenetic independent contrasts (PIC). The analysis was based on 55 species for which dynamic responses to temperature were quantified (one outlier excluded). Pearson's correlation coefficient ($r$) and its $p$-value are indicated in the figure.

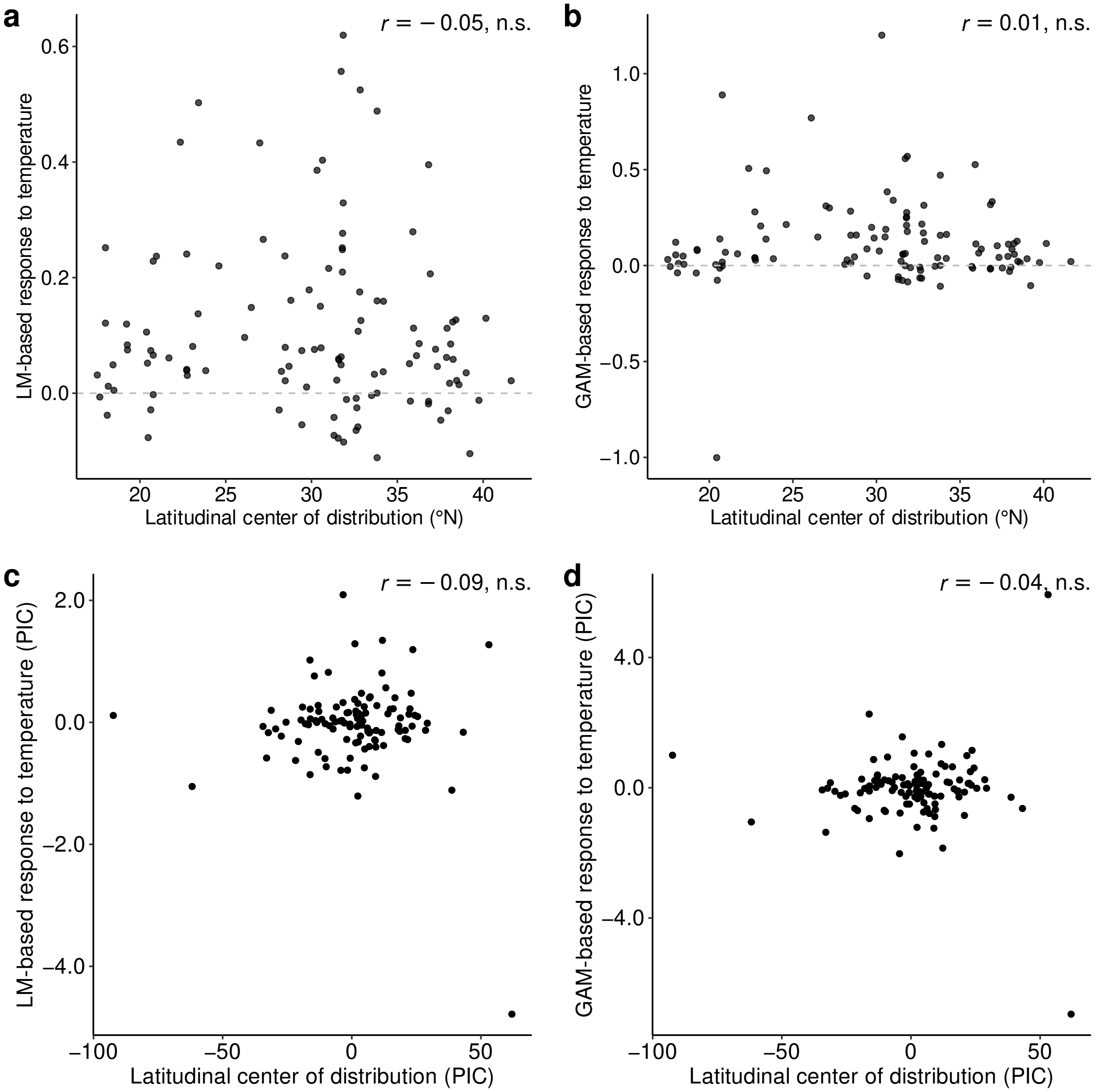


**Extended Data Figure 3| Relationships between latitudinal distribution and conventional temperature response traits.** Relationships between species' latitudinal distribution centers and their temperature responses estimated by linear model (LM) or generalized additive model (GAM). **a**, Latitudinal distribution center versus LM-based response (slope of the linear model between temperature and abundance). **b**, Latitudinal distribution center versus GAM-based response (mean slope of the first derivative of the fitted GAM). **c**–**d**, Corresponding relationships accounting for phylogenetic relatedness using phylogenetic independent contrasts (PIC). Pearson's correlation coefficient ($r$) and its $p$ value are indicated in the figure.

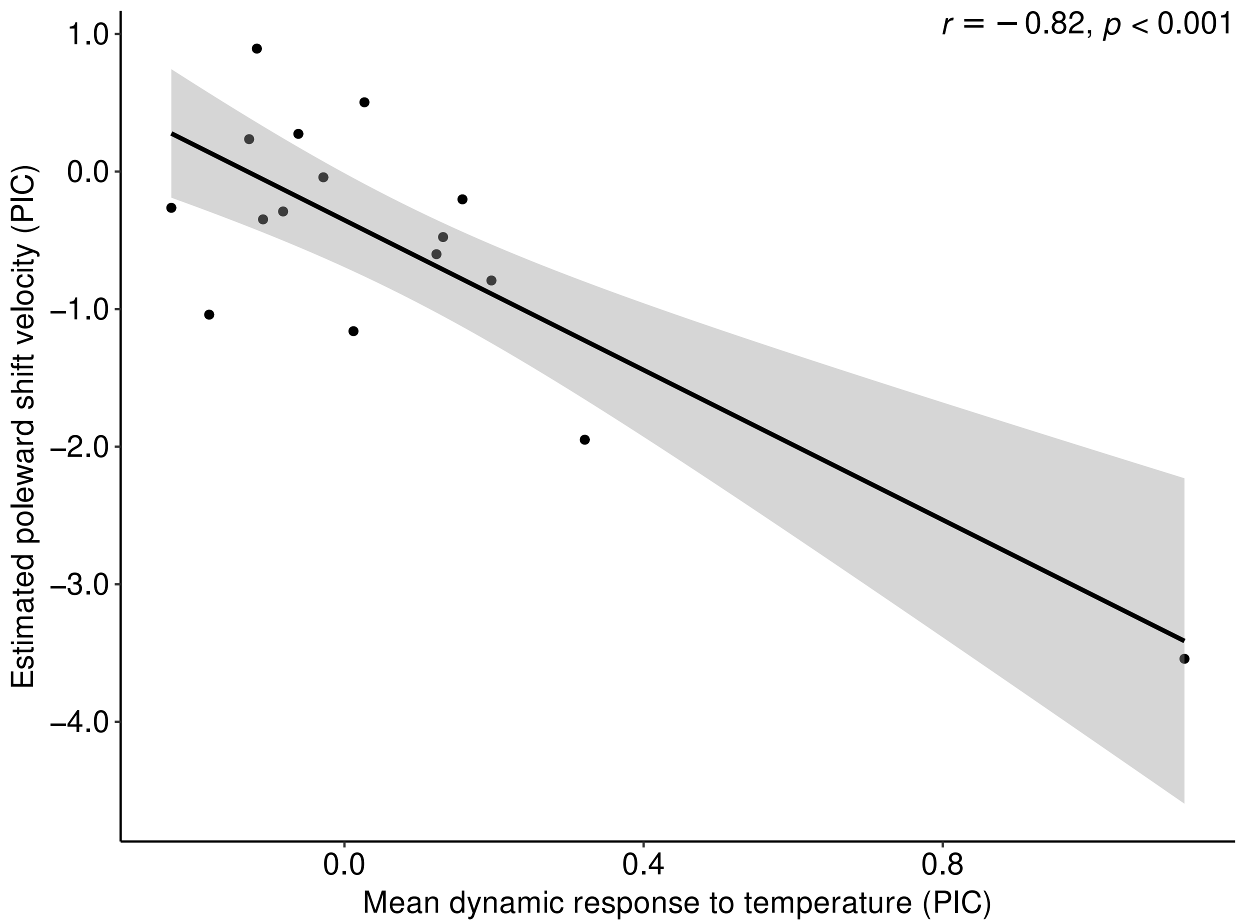


**Extended Data Figure 4| Relationship between fish range shift velocity and dynamic response to temperature accounting for phylogeny.** The relationship between each species' mean dynamic response to temperature and its estimated poleward shift velocity, based on iNaturalist and GBIF occurrence records collected in East Asia and Oceania, accounting for phylogenetic relatedness using phylogenetic independent contrasts (PIC), is shown. Pearson's correlation coefficient ($r$) and its $p$ value are shown.

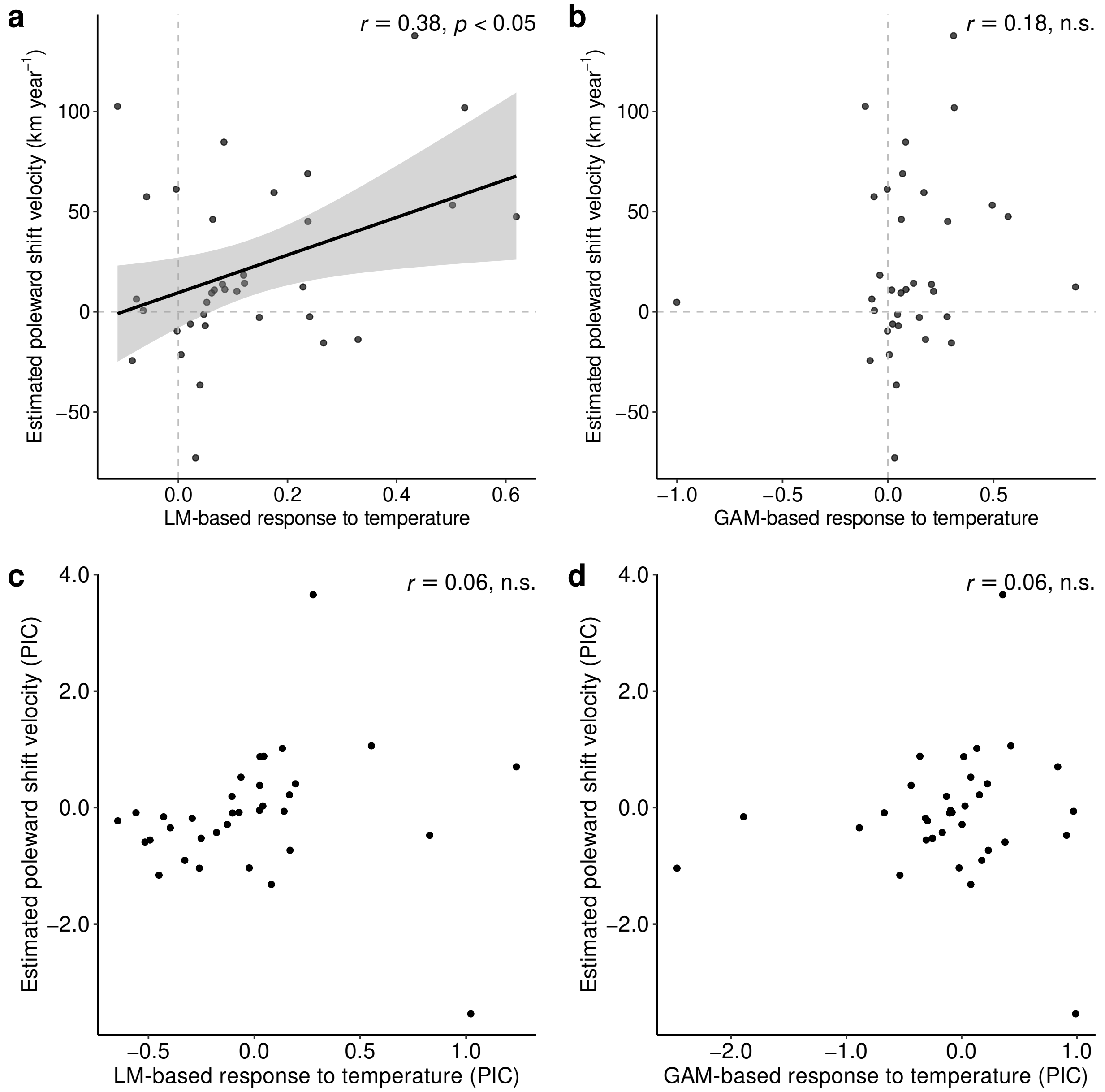


**Extended Data Figure 5| Relationships between conventional temperature response traits and fish range shift velocities.** Relationships between species' temperature response traits, quantified using linear model (LM) or generalized additive model (GAM), and their estimated poleward range shift velocities derived from iNaturalist and GBIF occurrence records collected in East Asia and Oceania, are shown. **a**, LM-based response (slope of the linear model between temperature and abundance) versus poleward shift velocity. **b**, GAM-based response (mean slope of the first derivative of the fitted GAM) versus poleward shift velocity. **c**–**d**, Corresponding relationships accounting for phylogenetic relatedness using phylogenetic independent contrasts (PIC). Pearson's correlation coefficient ($r$) and its $p$ value are shown.

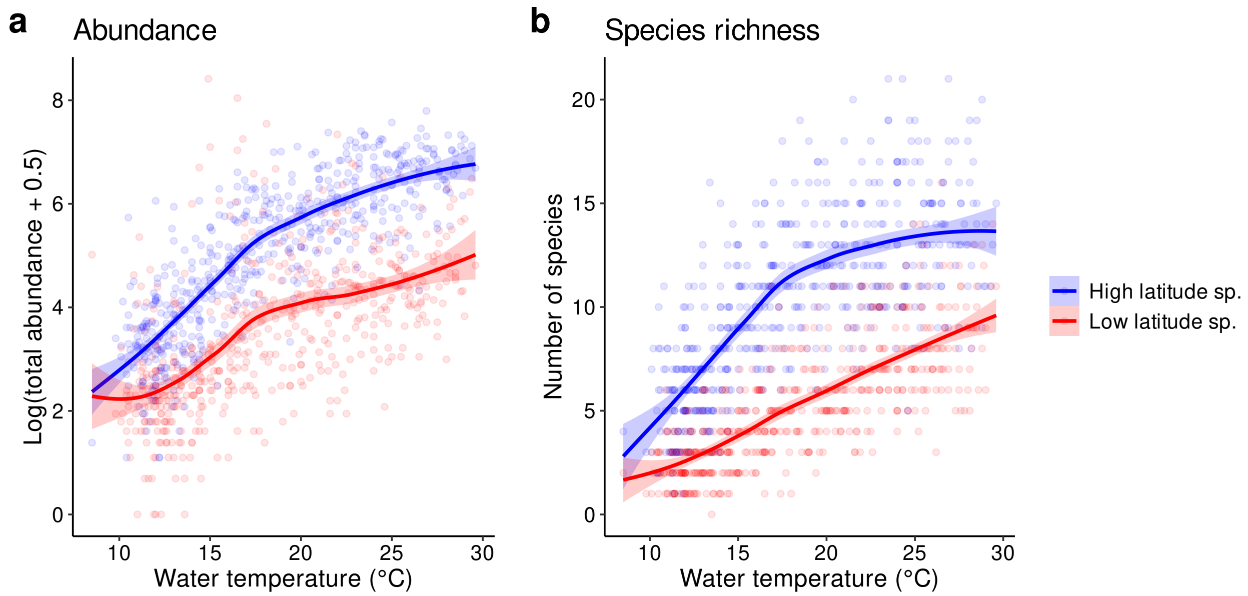


**Extended Data Figure 6| Relationships between water temperature and fish communities grouped by latitudinal category. a–b**, Fish species were categorized as "Low latitude" or "High latitude" depending on whether their latitudinal distribution centers were below or above the median of the 113 observed species. Relationships between water temperature and (**a**) total abundance and (**b**) species richness. Lines indicate smoothed fits using locally estimated scatterplot smoothing.

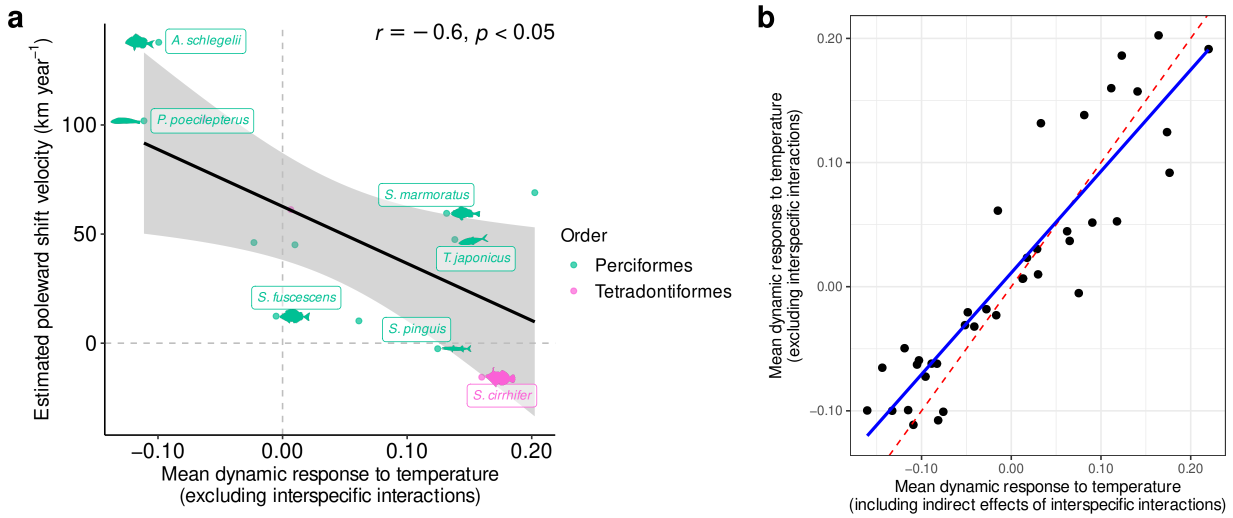


**Extended Data Figure 7| Dynamic responses to temperature after excluding interspecific interactions.** **a**, Relationship between the mean dynamic response to temperature after explicitly accounting for interspecific interactions (i.e., temperature responses without interspecific interactions) and species' estimated poleward range shift velocities derived from iNaturalist and GBIF occurrence records collected in East Asia and Oceania. Pearson's correlation coefficient ($r$) and its $p$ value are shown. **b**, Comparison between the mean dynamic responses to temperature estimated with and without accounting for interspecific interactions. The red dashed line indicates the 1:1 relationship, and the blue line shows the linear regression fitted to the data.

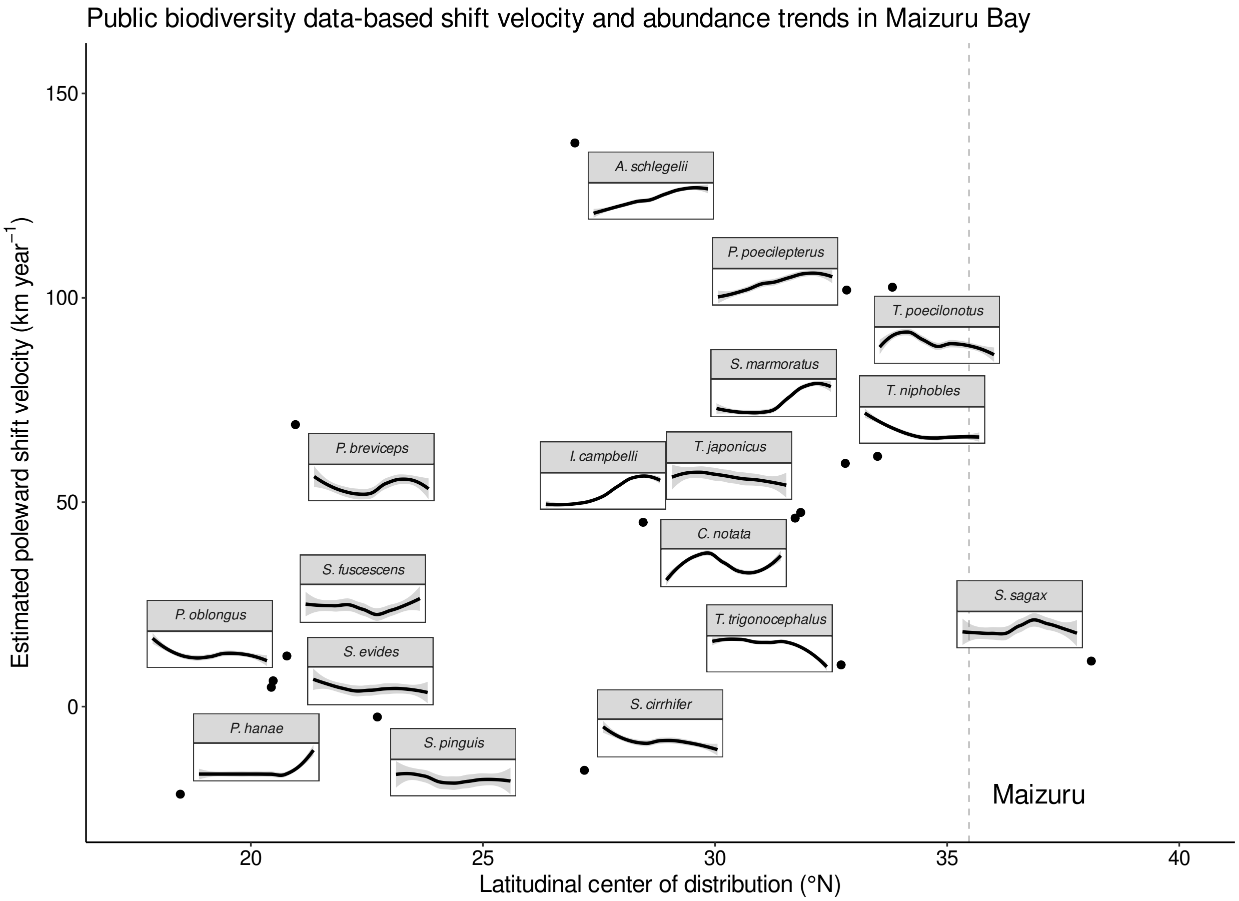


**Extended Data Figure 8| Relationships between species' latitudinal distribution centers and their estimated poleward shift velocities, with temporal changes in local abundance.** Each point represents a fish species, showing the relationship between its latitudinal distribution center and its estimated poleward shift velocity. Gray dashed line indicates the latitude of Maizuru Bay. For each species, its temporal trend in log-transformed abundance observed at the Maizuru site is shown as a small inset plot positioned at its corresponding point. Axes of the inset plots were removed for visualization purposes.
