## Supplementary Information for "Population-level, state-dependent response as a trait predicting species redistribution under climate change"

Contents:

**Supplementary Text**: Supplementary results and discussion:

1. Dynamic response to temperature after accounting for interspecific interactions,
2. Validation of the range shift velocity estimates from public biodiversity records using long-term local abundance trends,
3. Phylogenetically controlled correlations,
4. Interpretation of the dynamic responses to temperature of the top 5 positively and negatively responding species
5. Credits for fish silhouettes

Supplementary Text

1 ***Dynamic response to temperature after accounting for interspecific interactions***

The estimation of dynamic responses to temperature without considering effects from other species could be biased. Thus, we quantified dynamic responses to temperature after explicitly accounting for interspecific interactions and examined the robustness of the results.

First, to consider interspecific interactions, we selected 46 fish species that were observed in at least 5% of total sampling occasions (≥27 observations). We tested interspecific interactions among these species within the framework of UIC^1^, conditioning on water temperature, using the uic_across function in the *macam* package in R^2^. Then, for each focal species, we identified other species that showed significant causal influences on its population dynamics (including time lags up to −12, corresponding to a 6-month lag). These interspecific interactions, together with water temperature (including their time lags) were incorporated in the MDR S-map^3^. By explicitly including the effects of interacting species, we reconstructed the system’s state space and quantified the dynamic response to temperature after removing the effects of interspecific interactions.

Using this interspecific interaction-controlled framework, the mean dynamic response to temperature showed a slightly stronger association with species’ estimated poleward shift velocities (*r* = −0.6; Extended Data Fig. 7a) compared to estimates that did not explicitly account for interspecific interactions (*r* = −0.52; Fig. 4a). This improvement likely reflects a more accurate quantification of temperature-driven responses after disentangling the confounding effects of species interactions.

The absolute magnitude of mean dynamic responses was generally reduced when interspecific interactions were explicitly included in the model (Extended Data Fig. 7b), suggesting that part of the response previously attributed to temperature alone captured the effects of biotic interactions. Importantly, however, temperature responses estimated with and without accounting for interspecific interactions closely followed the 1:1 relationship and exhibited highly consistent overall patterns (Extended Data Fig. 7b). Given that obtaining time series for all interacting species is often impractical in real-world application, this consistency indicates that the main conclusions are robust even when interspecific interactions cannot be explicitly modeled.

2 ***Validation of the range shift velocity estimates from public biodiversity records using long-term local abundance trends***

To validate the estimated poleward range shift velocities derived from iNaturalist and GBIF records, we compared them with long-term monitoring data from Maizuru Bay spanning over 22 years. Extended Data Fig. 8 demonstrates that the range shift estimates are broadly consistent with observed population trends at the fixed site. Species whose literature-derived distribution centers^4,5^ are located relatively south of Maizuru but exhibit high estimated shift velocities—such as *Acanthopagrus schlegelii*, *Parajulis poecilepterus*, *Sebastiscus marmoratus*, and *Istigobius campbelli*—showed general increases in abundance during the monitoring period. Conversely, species whose literature-derived distribution centers are relatively close to Maizuru yet exhibit large poleward shift velocities—such as *Takifugu poecilonotus* and *Takifugu niphobles*—showed consistent declines in abundance, suggesting that their current realized distributional centers may have already shifted northward beyond Maizuru. These patterns indicate that the range shift velocities estimated from public biodiversity records reliably reflect realized species’ range shifts.

3 ***Phylogenetically controlled correlations***

To confirm that the observed correlations were not driven by phylogenetic relatedness among species, we repeated the correlation tests using phylogenetically independent contrasts (PIC) based on the reconstructed tree described in Methods. The negative correlations between species’ dynamic responses to temperature and both their latitudinal distribution centers and estimated poleward shift velocities remained significant even after accounting for phylogeny (Extended Data Fig. 2, 4). In contrast, correlations obtained using conventional LM- or GAM-based response traits were not significant when phylogenetic effects were considered (Extended Data Fig. 3c–d, 5c–d). Notably, the LM-based response trait even showed a positive correlation with poleward shift velocity when phylogeny was not accounted for (Extended Data Fig. 5a), but this relationship disappeared after phylogenetic correction (Extended Data Fig. 5c). This indicates that ignoring phylogeny can lead to misleading associations among traits, whereas our dynamic response trait robustly reflects species-specific ecological characteristics independent of phylogenetic conservatism.

4 ***Interpretation of the dynamic responses to temperature of the top 5 positively and negatively responding species***

Here, we provide ecological interpretations of the dynamic responses to temperature for the five species showing the strongest positive responses and the five species showing the strongest negative responses.

4.1 **Positive responders**
    4.1.1 *Epinephelus awoara* (mean dynamic response = 0.22)

Most species of *Epinephelus* (groupers) are distributed from tropical to warm-temperate waters^5^. *Epinephelus awoara* and *E. akaara* occur at relatively high latitudes but are still of warm-water origin. Catches of these species in set nets located in the Maizuru area tend to be high in autumn, implying southward migration as temperature decreases (Masuda et al., unpublished).

4.1.2 *Sillago japonica* (mean dynamic response = 0.18)

This species is carnivorous and actively feeds on benthic prey during the warm season^6^.

4.1.3 *Sphyraena pinguis* (mean dynamic response = 0.17)

This species is a piscivorous, pelagic, migratory fish. Individuals found in the survey area are mostly juveniles that form shoals, sometimes with other species such as the jack mackerel *Trachurus japonicus*.

4.1.4 *Petroscirtes breviceps* (mean dynamic response = 0.16)

This blenny is often found associated with drifting algae. Both juveniles and adults occur in the survey area. The species is distributed from tropical to temperate waters, and our survey area is close to the northern limit of their distribution.

4.1.5 *Epinephelus akaara* (mean dynamic response = 0.14)

This species is mentioned in Section 4.1.1.

4.2 **Negative responders**
    4.2.1 *Ditrema temminckii* (mean dynamic response = −0.16)

This species is viviparous and reported to mate from September to December^7^. As with other surfperches, it dominates in cold-temperate waters.

4.2.2 *Pseudolabrus sieboldi* (mean dynamic response = −0.14)

Unlike other wrasse species in the survey area, *P. sieboldi* tends to be solitary and rarely forms shoals. It is active at relatively low water temperatures.

4.2.3 *Hexagrammos otakii* (mean dynamic response = −0.13)

Maturation of the greenling *Hexagrammos otakii* is triggered when water temperature declines below 18 °C^8^. Therefore, increasing water temperature is likely to shorten the spawning season of this species. After spawning, males of *H. otakii* guard fertilized eggs until hatching. Egg-guarding males were common until 2004 but have not been observed since.

4.2.4 *Acentrogobius virgatulus* (mean dynamic response = −0.12)

This species is a benthic goby commensal with alpheid shrimp that construct burrows. It was formerly very abundant in the survey area when the census began, but its abundance has declined in recent years. This decline may be partly due to competition with the warm-water goby *Istigobius campbelli*, which has increased more recently^9^.

4.2.5 *Acanthopagrus schlegelii* (mean dynamic response = −0.11)

Black sea bream *Acanthopagrus schlegelii* is known to spawn in coastal waters and often migrates into estuaries and rivers. Most individuals found in the survey area are adults with an average body length of approximately 30 cm^10^. The species used to be absent in mid-winter up until 2016 but is now present year-round. It is currently one of the few species observed during the coldest season in the survey area.

5 ***Credits for fish silhouettes***

All fish silhouettes used in the figures were obtained from PhyloPic and are reproduced under their respective licenses. When silhouettes of the focal species were unavailable, silhouettes of morphologically similar species or higher taxonomic groups were used as proxies for visualization purposes only.

- *Engraulis japonicus*: *Engraulis ringens* by Juan Carlos Jerí (CC0 1.0 Universal Public Domain Dedication)
- *Sebastes cheni*: *Sebastes diaconus* by Felix Vaux (CC0 1.0 Universal Public Domain Dedication)
- *Trachurus japonicus*: *Scomberoides lysan* by Richard J. Harris (Public Domain Mark 1.0)
- *Pterogobius zonoleucus*: *Lythrypnus dalli* by tmccraney (CC0 1.0 Universal Public Domain Dedication)
- *Tridentiger trigonocephalus*: *Gobius* by tmccraney (CC0 1.0 Universal Public Domain Dedication)
- *Sphyraena pinguis*: *Sphyraena* by Erika Schumacher (Attribution-NonCommercial 3.0 Unported)
- *Paralichthys olivaceus*: *Paralichthys californicus* by Nick Schooler (Public Domain Mark 1.0)
- *Stephanolepis cirrhifer*: *Monacanthus ciliatus* by Emily Troyer (Attribution 4.0 International)
- *Parajulis poecilepterus*: *Coris julis* by Cesc Gordó-Vilaseca (CC0 1.0 Universal Public Domain Dedication)
- *Acanthopagrus schlegelii*: *Stenotomus chrysops* by Nathan Hermann (CC0 1.0 Universal Public Domain Dedication)
- *Sardinops sagax*: *Clupea harengus* by Nathan Hermann (CC0 1.0 Universal Public Domain Dedication)
- *Sebastiscus marmoratus*: *Sebastes fasciatus* by Nathan Hermann (CC0 1.0 Universal Public Domain Dedication)
- *Siganus fuscescens*: *Siganus rivulatus* by perevolotsky (CC0 1.0 Universal Public Domain Dedication)
